# Temporal discounting correlates with directed exploration but not with random exploration

**DOI:** 10.1101/783498

**Authors:** Hashem Sadeghiyeh, Siyu Wang, Maxwell R. Alberhasky, Hannah M. Kyllo, Amitai Shenhav, Robert C. Wilson

## Abstract

The explore-exploit dilemma describes the trade off that occurs any time we must choose between exploring unknown options and exploiting options we know well. Implicit in this trade off is how we value future rewards — exploiting is usually better in the short term, but in the longer term the benefits of exploration can be huge. Thus, in theory there should be a tight connection between how much people value future rewards, i.e. how much they discount future rewards relative to immediate rewards, and how likely they are to explore, with less ‘temporal discounting’ associated with more exploration. By measuring individual differences in temporal discounting and correlating them with explore-exploit behavior, we tested whether this theoretical prediction holds in practice. We used the 27-item Delay-Discounting Questionnaire to estimate temporal discounting and the Horizon Task to quantify two strategies of explore-exploit behavior: directed exploration, where information drives exploration by choice, and random exploration, where behavioral variability drives exploration by chance. We find a clear correlation between temporal discounting and directed exploration, with more temporal discounting leading to less directed exploration. Conversely, we find no relationship between temporal discounting and random exploration. Unexpectedly, we find that the relationship with directed exploration appears to be driven by a correlation between temporal discounting and uncertainty seeking at short time horizons, rather than information seeking at long horizons. Taken together our results suggest a nuanced relationship between temporal discounting and explore-exploit behavior that may be mediated by multiple factors.

## Introduction

The explore-exploit dilemma refers to a ubiquitous problem in reinforcement learning in which an agent has to decide between exploiting options it knows to be good and exploring options whose rewards are unknown [1]. For example, when ordering sushi at a favorite restaurant, should we exploit our usual favorite (the Rainbow Roll), which is guaranteed to be good, or explore the Burrito Roll, which could be delicious, disgusting or somewhere in between. As anyone who has agonized over a dining decision will know, making explore-exploit choices can be hard, and there is considerable interest in how these decisions are made by humans and other animals [2].

Recently, a number of studies have shown that people make explore-exploit decisions using a mixture of two strategies: directed exploration and random exploration [3–8]. In directed exploration, choices are biased towards more informative options by an ‘information bonus,’ that increases the relative value of unknown options [9]. In random exploration, behavioral variability, perhaps driven by random noise processes in the brain, causes exploratory options to be chosen by chance [1, 10]. Further work suggests these two types of exploration have different computational properties [4], age dependence [11], and may be controlled by different systems in the brain [12–15].

Regardless of the type of exploration, the benefits of exploring over exploiting lie in the possibility of earning larger rewards in the future. For example, in our restaurant example, if the Rainbow Roll is an above-average item on the menu, then, in the short term, exploiting it will usually be best. In the longer term, however, if the Burrito Roll turns out to be sublime, then we could order this roll again and again for years to come. Thus, how much we care about future rewards, that is how we discount them relative to immediate rewards, should play a critical role in how we make our explore-exploit choice.

Optimal models of explore-exploit decision making formalize this relationship between temporal discounting and exploration, at least for directed exploration [9]. In these models, the explore-exploit choice is made by choosing the option that maximizes the expected discounted future reward. Because this maximizing behavior is deterministic (apart from rare cases in which options are tied), optimal models do not exhibit random exploration. Thus, while they predict a negative relationship between temporal discounting and directed exploration, they say nothing about the relationship with random exploration. Sub-optimal models of explore-exploit decision making do include random exploration, but most of them predict no relationship with temporal discounting [1, 10, 16].

Thus, in theory, one might predict a negative relationship between temporal discounting and directed exploration, and no relationship between temporal discounting and random exploration. In practice, however, previous experimental work suggests a more nuanced picture because of how temporal discounting covaries with our attitudes toward risk. In particular, high temporal discounting is associated with greater impulsivity [17], and higher impulsivity is associated with greater risk taking [18]. This suggests that more temporal discounting is associated with more risk seeking [19, 20] (However, by defining risk seeking in terms of probability discounting, some studies on the relationship between temporal and probability discounting have yielded ambiguous results on this suggestion [21–25]). In most explore-exploit paradigms, such increased risk taking would look a lot like increased directed exploration, because the more informative option is usually more uncertain, i.e. risky, too. Thus, while theory might predict a negative correlation between temporal discounting and directed exploration, this effect could be countered by a positive correlation between temporal discounting and risk taking.

In the current study, we investigated the correlation between temporal discounting and the two kinds of exploration using an individual differences approach. That is, we asked whether people with higher temporal discounting show less directed and/or random exploration. We used the 27-item Delay Discounting Questionnaire [26] to measure temporal discounting. In this questionnaire, participants choose between between small but immediate amounts of money and a larger but delayed amounts of money (e.g. $11 now or $30 in two weeks). Based on participants’ pattern of choosing between immediate and delayed options, a parameter *k* [27] is calculated for each participant which estimates their average discounting rate for delayed rewards.

We used the Horizon Task [3] to measure directed and random exploration. In this task participants make a series of choices between two slot machines (one-armed bandits). When played, each machine pays out a reward from a Gaussian distribution. The average payout is different for each machine such that one option is always better on average. Thus, to maximize their rewards, participants need to exploit the option with the highest average payout, but can only find out which option is best by exploring both options first. By manipulating key parameters in this task (distribution of rewards, time horizon, and the amount of uncertainty for each bandit), the Horizon Task allows us to quantify directed and random exploration, and, crucially, to dissociate them from baseline risk seeking and behavioral variability.

Thus, by comparing individual differences in behavior on the Horizon Task with individual differences in temporal discounting, we aimed to quantify the relationship between the two types of exploration and temporal discounting.

## Methods

### Participants

We collected data from a total of 82 participants (ages 18-25, average = 19.10; Females = 47, Males = 35). Participants were recruited through the Psychology subject pool at the University of Arizona and received course credit for their participation. All participants gave informed consent and the study was approved by the Institutional Review Board at the University of Arizona and all experiments were performed in accordance with relevant guidelines and regulations.

### Temporal Discounting measure

To measure temporal discounting we used the Delay Discounting Questionnaire developed by [28]. In this instrument there are 27 questions asking participants’ preferences between two hypothetical monetary rewards: one of which pays immediately but is smaller, and the other pays more but is delayed. For example, one item asks: Do you prefer $11 today or $30 in 7 days? The amount of smaller-immediate reward (“today” option), larger-delayed reward (“later” option) and the delay (in terms of days) vary in those 27 questions (“today” reward between $11 - $80; “later” reward between $25 - $85; Delay between 7 - 186 days). The exact values are reported in [28]-Table 3.

One out of four participants were selected by chance (by drawing a card at the end of experiment) to receive the actual money according to their responses. If a participant drew a winning card (%25 chance), they then would proceed to draw a numbered chip from a bag (out of 27 chips numbered from 1 to 27 according to the number of items in the monetary choice questionnaire). The number on the chip corresponds to the number of the question we would look at for the actual pay-out. For example, if the winning participant picked the number 19 and they answered “later” on the question #19: “Do you prefer $33 now or $80 in 14 days?”, they need to come back to lab in 14 days and receive $80 in cash after signing a receipt form.

To quantify temporal discounting we used a number of different measures. The simplest was just the number of today options chosen, with greater temporal discounting associated with larger number of “today” choices.

More sophisticated measures of temporal discounting were obtained by fitting a hyperbolic discount factor to the data. In particular, we assume that future reward, *A*, arriving after a delay *D*, is discounted according to a hyperbolic discount factor [29]:

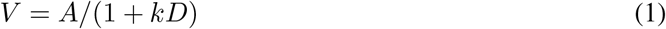

where *k* is the subject-specific discount factor. Fitting *k* was done using the spreadsheet provided by [30] based on the method described in [27]. In addition to computing an overall *k* using all 27 items, this approach also computes separate discount factors for small, medium and large reward items, based on the idea that delay discounting may be different for different range of rewards, and also the geometric mean of the small, medium and large *k*s. Based on the range of monetary values, the 27 choices are divided into three 9-item categories: small, medium and large ranges. Then, based on the hyperbolic discounting equation (Eq. 1), it finds a k value for each item as a point in which there is no difference between choosing “today” and “later” options for that item. Then for each participant based on his/her answers and the patterns of switches from “today” to “later” options and the reverse, it gives us a k-value for each 9-item category: Small k, Medium k, Large k.

For example, in question 2 it asks: Would you prefer $55 today, or $75 in 61 days? The indifference point is when the $75 in 61 days worth as $55 today. We can calculate the k for the indifference point, in which the “today” and “later” choices look the same, by plugging *V* = 55, *A* = 75, *D* = 61 in Eq. (1):

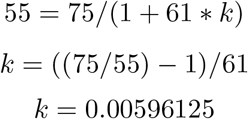

If a participant choose “today” for this question, they have a *k* > 0.00596125.

Similarly, if the same participant answer “later” in question 7: Would you prefer $15 today, or $35 in 13 days?, the indifference point would be *k* = ((35/15) − 1)/13 = 0.102564103 so our participant would have a *k* < 0.102564103. So for this participant given these two questions, we can estimate their k to be between 0.00596125 < *k* < 0.102564103. By adding more questions, we can obtain better estimates for *k*.

Thus we have six measures of temporal discounting for each subject: the fraction of “today” choices, overall *k*, small *k*, medium *k*, large *k*, and the geometric mean of small, medium and large *k*s.

### Horizon Task

The Horizon Task [3] is a recently developed task that allows for the measurement of directed and random exploration. The key manipulation in the Horizon Task is the time horizon, the number of trials participants will make in the future. The idea being, that in a long time horizon, people should explore, while in a short time horizon, people should exploit. Thus the change in behavior between short and long horizons can be used to quantify directed and random exploration.

More specifically, in the Horizon Task participants choose between two one-armed bandits. When chosen, the bandits pay out rewards sampled from a Gaussian distribution whose standard deviation is always fixed at 8 points, but whose mean is different for each bandit and can change from game to game. Each game lasts for 5 or 10 trials and participants’ job is to make multiple choices between the two bandits to try to maximize their reward. Because they know nothing about the mean of each bandit at the start of each game, they can only find out which option is best by exploring.

To control the amount of information, the first four trials of each game are predetermined (Figure 1-B). Participants are instructed to pick either the left or right bandit during these four “forced trials”. By changing the number of forced choices for each bandit, we manipulate the amount of “uncertainty” or information participants have about the payoffs from each bandit. In the unequal uncertainty (or [1 3] condition) participants are forced to choose one option once and the other three times; whereas in the equal uncertainty (or [2 2] condition) participants play both options twice. After the forced-choice trials, the rest of trials are “free trials” in which participants make their own choice. The number of free trials varies between horizon conditions with 1 free choice in the horizon 1 condition and 6 free choices in the horizon 6 condition.

**Figure 1:**
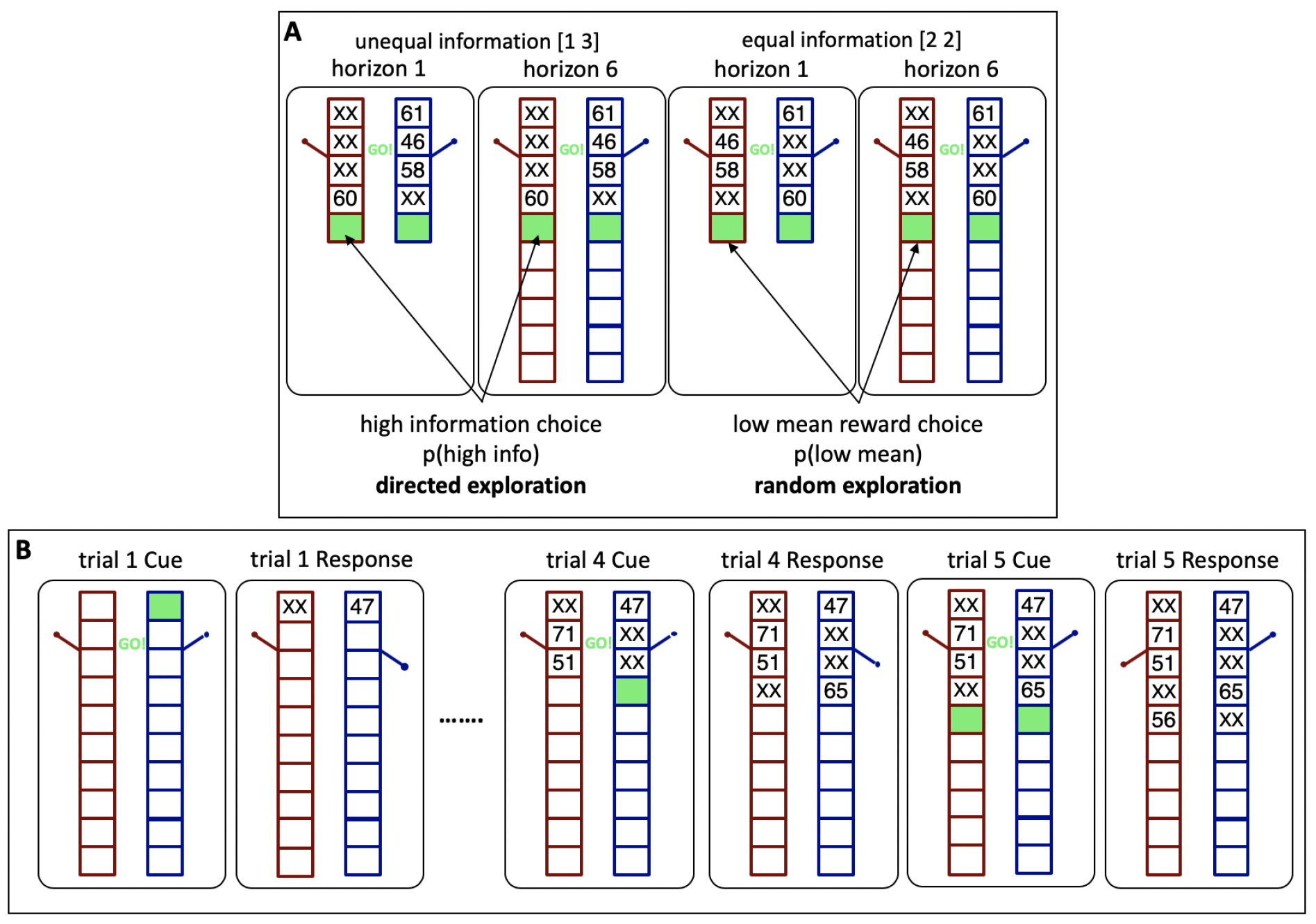
A) Horizon task: the four forced trials set up one of two information conditions (unequal [1 3] and equal [2 2] information) and two horizon conditions (1 vs 6) before participants make their first free choice. B) Active version of the horizon task

These two information conditions allow us to quantify directed and random exploration by looking at the first free choice in each game, immediately after the four forced choices (Figure 1-A). Because directed exploration involves information seeking, it can be quantified as the probability of choosing the more informative option in the [1 3] condition, p(high info). Conversely, because random exploration involves decision noise, it correlates with choosing the low mean option in the [2 2] condition, p(low mean). Computing these measures separately for each horizon condition allows us to quantify four key properties of explore-exploit behavior:

- uncertainty preference as p(high info) in horizon 1
- baseline behavioral variability as p(low mean) in horizon 1
- directed exploration as Δp(high info), the change in information seeking with horizon
- random exploration as Δp(low mean), the change in variability with horizon

In Supplementary Materials-4 you can find the onscreen instructions used to instruct participants at the beginning of the Horizon Task.

### Model-based Analysis

In addition to the above-mentioned model-free parameters (p(high info) and p(low mean)) we also fit a logistic model that was previously shown to be adequate in capturing the basics of the Horizon Task [3]. With this model we estimate two main parameters: “information bonus” and “decision noise” which corresponds to the model-free measures of directed and random exploration, respectively. The description of the model is provided in the Supplementary Materials-1. The modeling will help us to disentangle directed and random exploration more clearly. However, since there was a high correlation between model-free and model-based parameters (Supplementary Materials-2 Figure S1) and both the model-based and model-free parameters yielded the same relationships with the temporal discounting (Supplementary Materials-2 Figure S2), and since the model-free approach requires less assumptions than the model-based approach, we chose to include the model-free analysis in the main article and move the modeling part to the Supplementary Materials.

### Statistical Analysis

To evaluate the basic behavior on the Horizon Task, we used the paired (dependent) sample t-test. For directed exploration we looked to see whether there was a significant increase in the mean of p(high info) from horizon 1 to horizon 6 condition using the paired sample t-test. Similarly, for the random exploration we used paired sample t-test to see whether there was a significant increase in the p(low mean) between horizon 1 and horizon 6 condition.

To evaluate the relationship between measures of temporal discounting and the Horizon Task parameters, we simply calculated the Pearson correlation coefficients between the 6 measures of temporal discounting (the 5 k’s: overall k, small k, medium k, large k, geometric k and the total number of today items chosen) on one hand and the Horizon Task parameters (directed exploration, random exploration, p(high info) in horizon 1 and 6, p(low mean) in horizon 1 and 6, reaction time on the first free trial in horizon 1 and 6, and accuracy in horizon 1 and 6) on the other hand. Accuracy is defined as choosing the high mean option.

## Results

### Behavior on the Horizon Task (Model-free)

Table 1 shows the Mean and Standard Deviation (SD) for the basic task parameters in the sample. Figure 2 shows the distribution of basic task parameters in the sample (N = 82). Behavior on the Horizon Task was consistent with that previously reported in [3]. Specifically we see a significant increase in p(low mean) with horizon (p(low mean)h1_average = 0.2883; p(low mean)h6_average = 0.3554; t(81) = 3.87; p < 0.001) (Figure 4-B) and we see a clear trend (but not significant) in p(high info) with horizon (p(high info)h1_average = 0.5146; p(high info)h6_average = 0.5486; t(81) = 1.75; p = 0.084) (Figure 4-A), consistent with participants using both types of exploration in this paradigm. Figure 3 shows the scatter plots comparing p(high info) and p(low mean) for individual participants in horizon 1 and horizon 6 conditions. Out of 82 participants, 57 showed random exploration (p(low mean) h6 > p(low mean) h1) and 47 showed directed exploration (p(high info) h6 > p(high info) h1) on average.

**Table 1:**
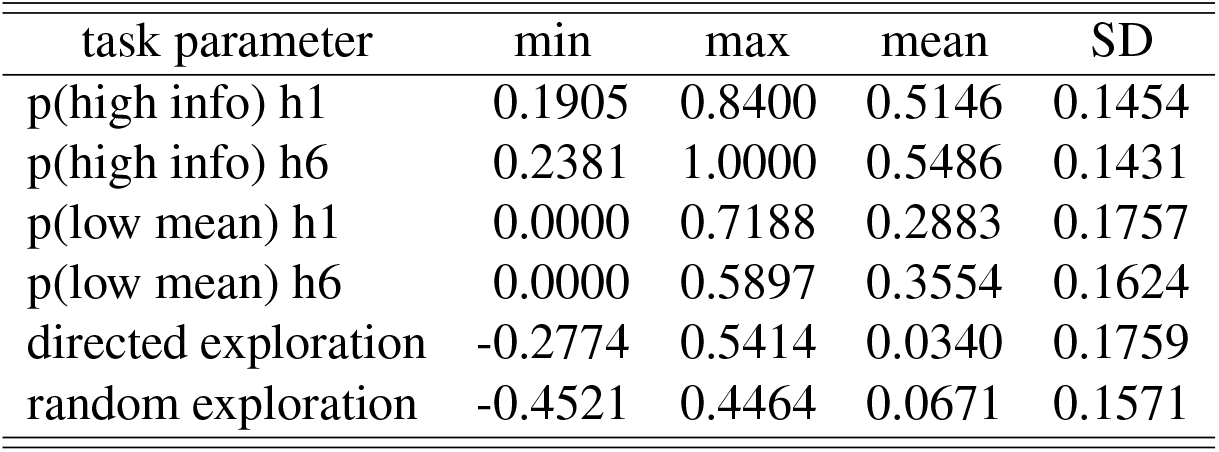
Ranges, Means and Standard Deviations for basic task parameters

**Figure 2:**
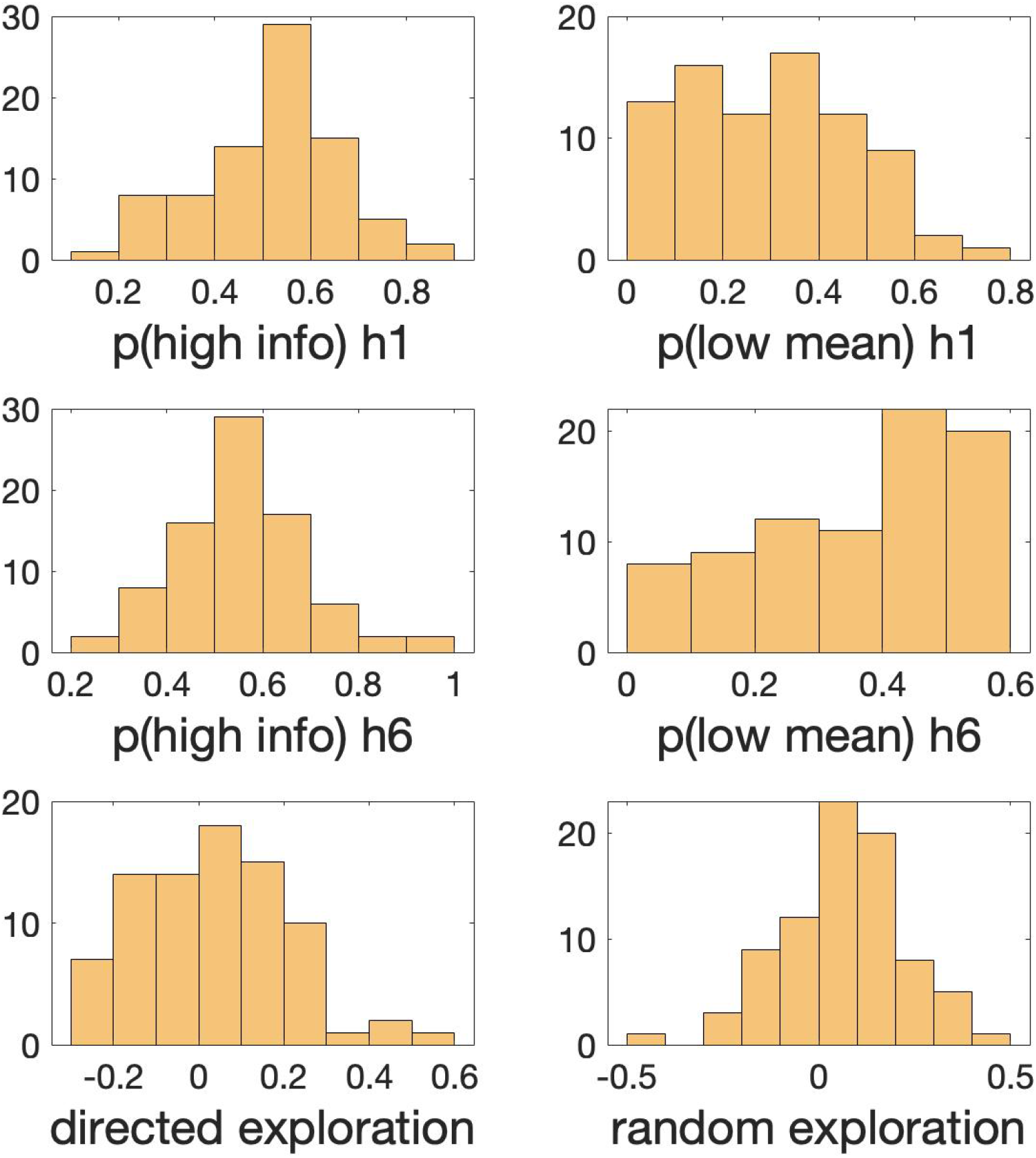
Histograms demonstrating the distribution of Horizon Task parameters (p(high info) and p(low mean) in horizon 1 & 6 and directed & random explorations) in our sample of 82 participants. The y-axis is the frequency or the number of occurrences per each value on the x-axis.

**Figure 3:**
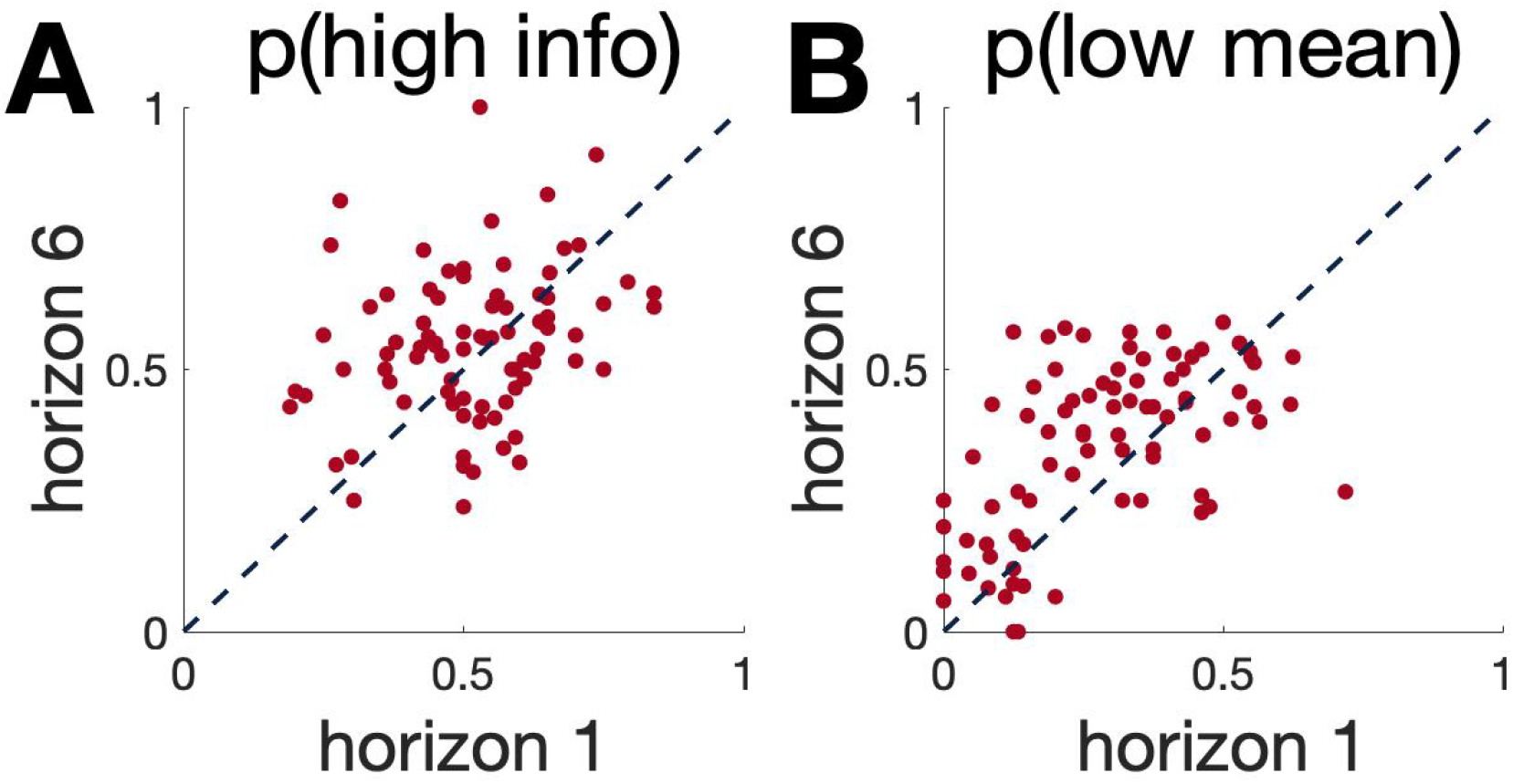
Scatter plots comparing task parameters (A) p(high info) and (B) p(low mean) for individual participants in horizon 1 and horizon 6. The dashed lines show equality. Those cases above this line denotes the expected horizon behavior (where p(high info) h6 > p(high info) h1 and p(low mean) h6 > p(low mean) h1.

**Figure 4:**
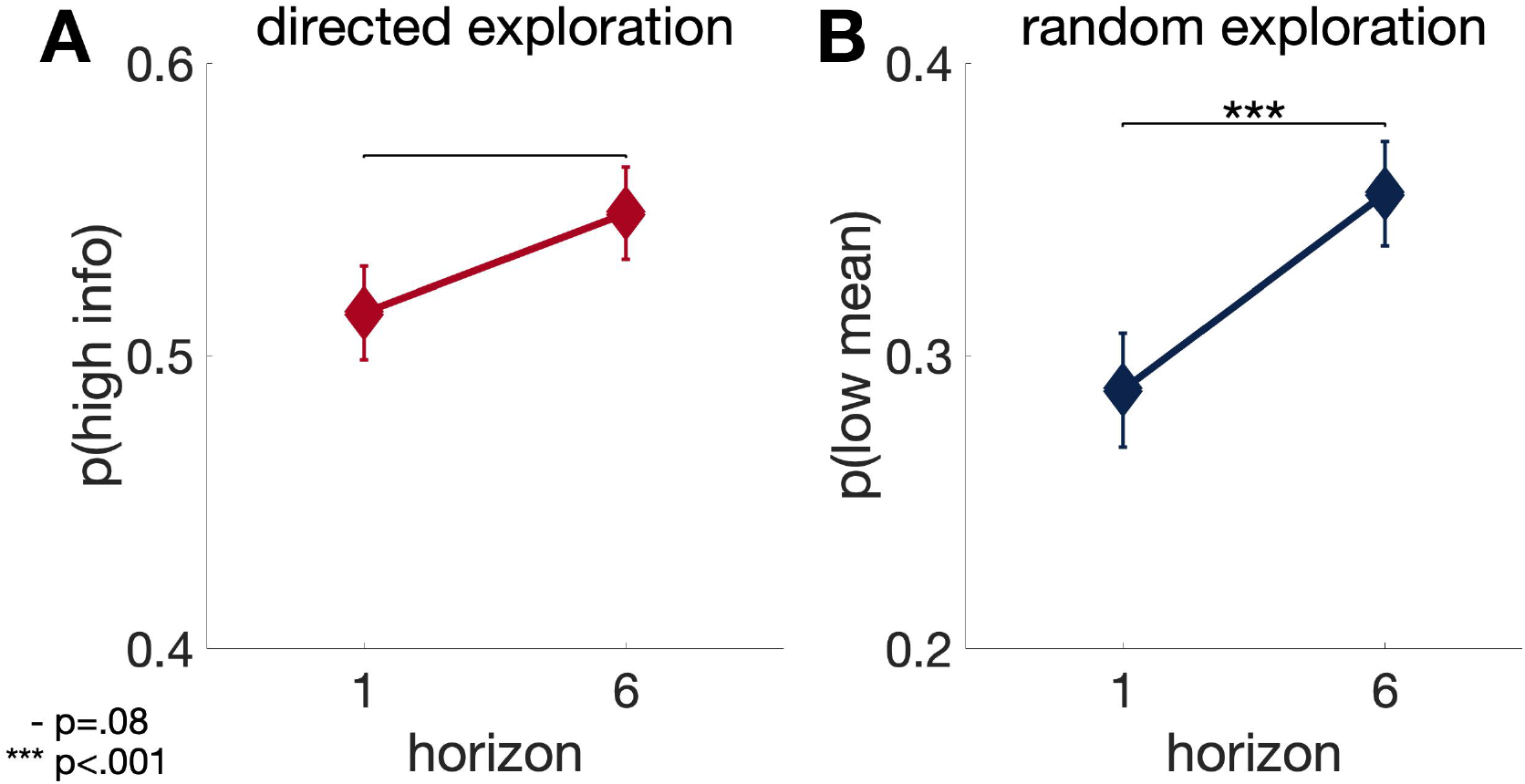
The average of p(high info) (A) and p(low mean) (B) for 82 participants on each horizon condition. The increase in p(high info) and p(low mean) from horizon 1 to horizon 6 follows the typical pattern observed in our previous studies and shows the use of both directed and random exploration.

### Behavior on the Temporal Discounting Task

For the temporal discounting measure we obtained 5 different *k* values for each participant as a measure of how much they discount future reward. We also can simply estimate that measure just by counting the number of times participants chose the immediate versus delayed reward (Supplementary Materials-3). Table 2 shows the range, mean and standard deviations of temporal discounting indices (*k*’s and # today items) in 82 participants of our study which is similar to previous studies using the same measure [26, 28]. Figure 5 shows the histogram of distribution of temporal discounting indices in the sample (N = 82).

**Table 2:** Ranges, Means, and Standard Deviations for temporal discounting measures

|  | min | max | mean | SD |
| --- | --- | --- | --- | --- |
| Overall k | 0.0004 | 0.2494 | 0.0303 | 0.0503 |
| Small k | 0.0016 | 0.2468 | 0.0476 | 0.0613 |
| Medium k | 0.0002 | 0.25 | 0.0297 | 0.0440 |
| Large k | 0.0002 | 0.2488 | 0.0201 | 0.0371 |
| Geomean k | 0.0004 | 0.2485 | 0.0276 | 0.0411 |
| # today items | 4 | 27 | 15.85 | 4.32 |

**Figure 5:**
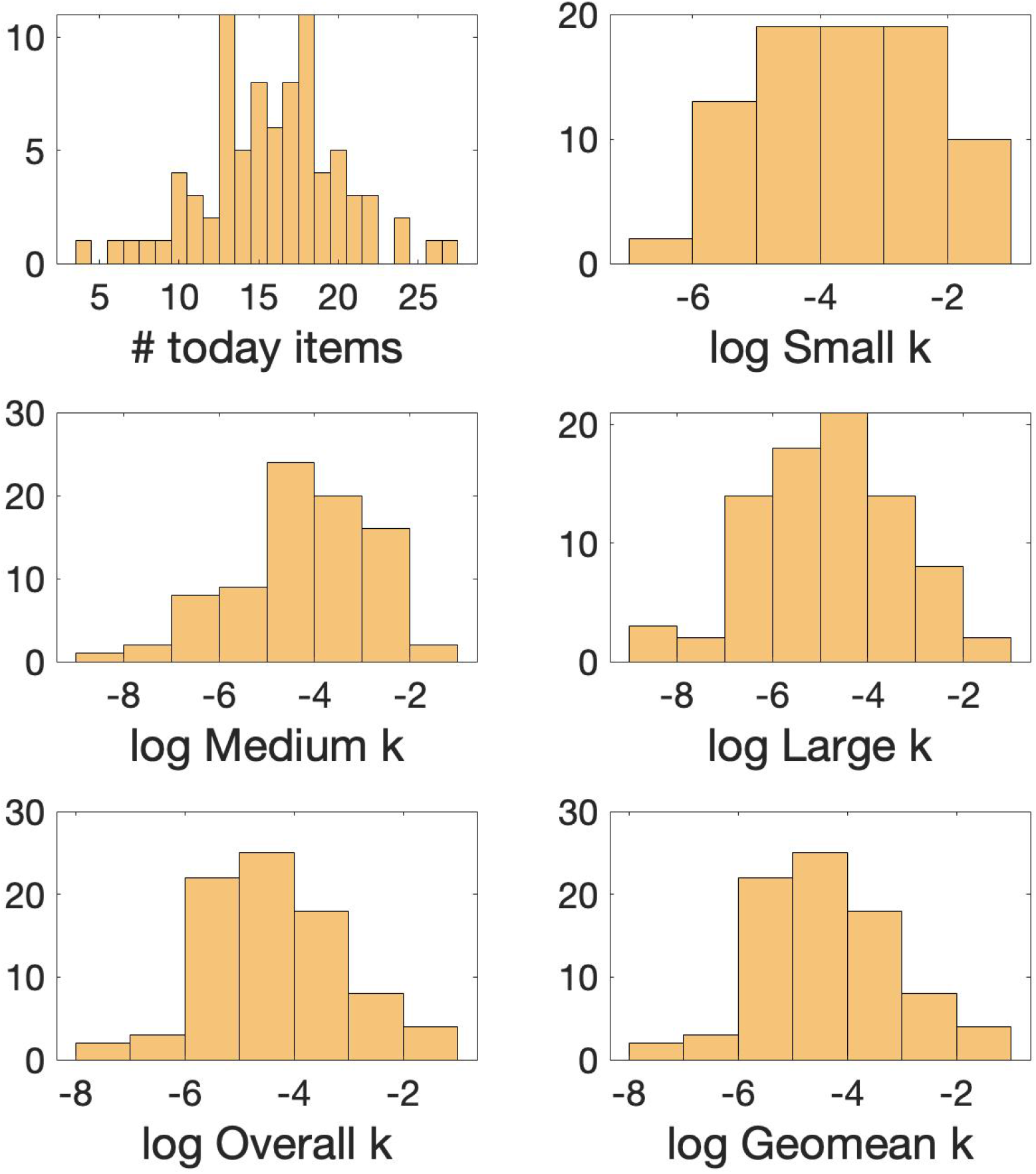
Histograms demonstrating the distribution of temporal discounting measures in our sample of 82 participants. The y-axis is the frequency or the number of occurrences per each value on the x-axis.

In our research, it turned out that all of these indices are highly correlated with each other (Supplementary Materials-3) and all have very similar relationship with directed and random exploration. The more simple measure of # today items has a Pearson’s correlation coefficient between .89 - 1 with the more complicated *k* measures (Supplementary Materials-3 Figure S3).

### Correlation between temporal discounting and explore-exploit behavior

Table 3 shows the correlation between measures of temporal discounting and the horizon task parameters: directed and random exploration, p(high info) & p(low mean) at horizons 1 & 6, reaction times and accuracy (the percentage of times the “accurate” option (the higher mean option) was chosen for each horizon (1 & 6) conditions. We found a significant negative correlation between between temporal discounting and directed exploration, with more temporal discounting associated with less directed exploration. Closer inspection revealed that this negative correlation was driven by a positive correlation between temporal discounting and p(high info) at horizon 1 and a zero correlation between temporal discounting and p(high info) at horizon 6.

**Table 3:** Correlation between task parameters and log (k overall)

|  | r | p |
| --- | --- | --- |
| directed exploration | **-0.30** | 0.007 |
| random exploration | 0.04 | 0.720 |
| p(high info) h1 | **0.35** | 0.001 |
| p(high info) h6 | -0.01 | 0.924 |
| p(low mean) h1 | 0.22 | 0.052 |
| p(low mean) h6 | **0.27** | 0.014 |
| accuracy h1 | **-0.33** | 0.003 |
| accuracy h6 | -0.17 | 0.130 |
| reaction time h1 | -0.05 | 0.669 |
| reaction time h6 | -0.12 | 0.304 |

In contrast to directed exploration, temporal discounting did not correlate with random exploration. There was, however, a positive correlation between temporal discounting and overall behavioral variability, p(low mean) in both horizon conditions. This suggests that people with higher temporal discounting perform worse on the task overall.

Finally, to demonstrate that the significant correlations were not driven by outliers, we plot the correlations between measures of directed and random exploration and the number of today items chosen in Figure 4.

### Model-based Analysis

We also utilized a logistic model (further explained in the Supplementary Materials-1) to estimated two main parameters“information bonus” and “decision noise” which are assumed to correspond to p(high info) and p(low mean) in the model-free analysis, respectively. Figure S1 in the Supplementary Materials-2 shows that in fact there are high correlations between model-free and model-based parameters. Additionally, Figure S2 shows that the correlations between temporal discounting and model-based parameters are similar to the correlations between temporal discounting and model-free parameters (Figure 6).

**Figure 6:**
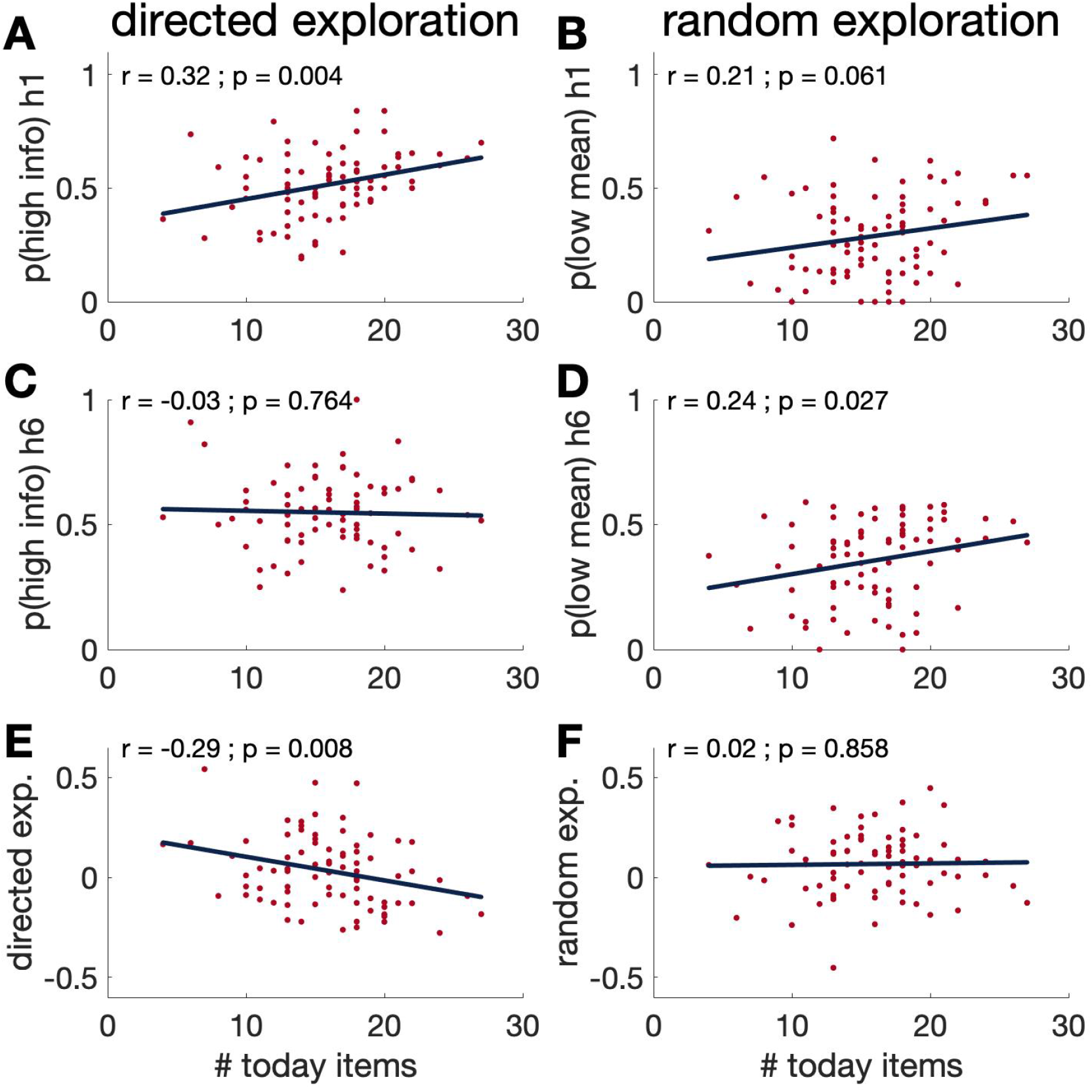
Scatter plots for (A) p(high info) h1, (B) p(low mean) h1, (C) p(high info) h6, (D) p(low mean) h6, (E) directed exploration and (F) random exploration over a temporal discounting measure (# today items). It clearly shows that the negative correlation between temporal discounting and directed exploration is driven by a positive correlation between temporal discounting and p(high info) h1.

## Discussion

In this study we investigated the correlation between temporal discounting measured by a monetary choice questionnaire [28] and two types of exploration (directed and random) measured by the Horizon Task [3]. We found a negative correlation between temporal discounting and directed exploration that was driven by a positive correlation between temporal discounting and uncertainty seeking in horizon 1. Conversely, we found no correlation between temporal discounting and random exploration, although we did see a positive correlation between temporal discounting and overall behavioral variability.

While the negative correlation between temporal discounting and directed exploration (i.e. Δ*p*(high info) is consistent with the theory, the correlation with *p*(high info) in each horizon condition is not. In particular, normative models predict a negative correlation between temporal discounting and *p*(high info) in horizon 6 and no correlation in horizon 1. Conversely, we found no correlation with horizon 6 behavior and a positive correlation with horizon 1 behavior.

One reason for this discrepancy could be the possible positive association between temporal discounting and risk taking [19, 20] (See [21–25] for suggesting otherwise). In both horizon conditions in the Horizon Task, the more informative option is also the more uncertain, riskier option. Thus, by this account, people who discount more would show greater *p*(high info) in both horizon conditions, but this would be counteracted by a negative relationship between temporal discounting and directed exploration in horizon 6. That is, in horizon 1, directed exploration is not present, and so the positive association with temporal discounting is revealed. In horizon 6, directed exploration is present, and this negative relationship with temporal discounting counteracts the positive relationship with risk taking leaving no correlation overall. Testing this hypothesis requires a future study that includes appropriate measures of risk taking.

The fact that random exploration does not correlate with temporal discounting is also consistent with theories of random exploration [1, 10]. Moreover, this apparent dissociation between directed and random exploration is consistent with other findings showing that directed and random exploration have different computational properties [4], different age dependence [11], and may rely on dissociable neural systems [12, 14, 15]. In this regard it is notable that directed exploration appears to rely on the same frontal systems thought to underlie temporal discounting [5, 12, 14, 31–33], while random exploration does not. Thus, an intriguing prediction is that the relationship between directed exploration and temporal discounting may be mediated by the integrity of frontal circuits, something that future neuroimaging studies could address.

There are several limitations in the current study. First, the chosen measures for both temporal discounting and exploratory behavior are very specific. This questions the generalizibility of our results. Although a strong correlation between different measures of temporal discounting has been demonstrated in several studies [34, 35], most of these measures are monetary which may have weak relationships with delay discounting in other domains [36]. Exploratory behavior also has been studied in different settings including foraging, repeated choice and sequential choice paradigms and it seems there is no shared factor underlying exploratory behavior in all of these tasks [37]. Replicating the current study using other measures of exploration and temporal discounting, will provide us with more evidence to better assess the generlizablilty of the current results.

Another important limitation of our study is recruiting university students as participants. Between all possible biases that such a selective sample may introduce in our study, age seems the most obvious one. It has been shown that temporal discounting [38], exploratory behavior [39] and risk-taking behavior [40], all varies significantly through the lifespan. So it is unclear how the results of the current study would look like in different age groups. This would be an interesting topic for a future study.

Lastly, we hypothesised the mediating role of risk taking to explain the results while we haven’t included appropriate scales to measure it in the current study. A future study can shed more light on this hypothesis by adding measures of risk taking.

## Acknowledgements

The authors thank Shlishaa Savita and Kathryn Lui Kellohen for their help in collecting and organizing data.

## Author Contributions Statements

HS, SW, AS and RCW designed the experiment. HS, MRA and HMK ran the experiment. HS and SW analyzed the data with supervision from RCW. HS and RCW wrote the manuscript with input from all other authors.

## Funding

The authors have no funding to disclose.

## Compliance with Ethical Standards

All procedures performed in experiments were in accordance with the ethical standards of the institutional research committee and with the 1964 Helsinki Declaration and its later amendments or comparable ethical standards.

## Conflicts of Interest

The authors declare that they have no conflict of interest.

## Informed Consent

Informed consent was obtained from all individual adult participants included in the study.

## Data Availability Statement

All the raw data and MATLAB codes for the analysis and plots are available at https://github.com/hashem20/temporal-discounting-explore-exploit

## Supplementary Materials

### 1- Model fitting

In addition to model-free parameters (p(high info) and p(low mean)), We used a simple logistic model to fit the decisions on the first free trial (trial 5). We hypothesized that the value of each option (a or b) is dependent on 3 main parameters: R, the average reward of the option (based on previous trials); I, the information, defined as whether choosing the option would provide you with valuable information so the less information we have about an option the more its information bonus; and s, spatial location, we hypothesized some subjects might be biased in choosing right over left option or vice versa. Therefore, we can express the value of each option as:

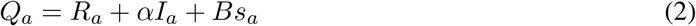

where *α* is called information bonus and B is the spatial bias.

We further assume the value of each option is perturbed with a logistic noise with the standard deviation of *σ*_*d*_. So the probability of choosing option a over b, assuming the participants have a linear utility function, will be calculated as:

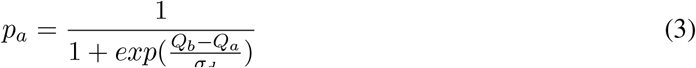

Replacing *Q*_*a*_ and *Q*_*b*_ with their equivalent from equation (2):

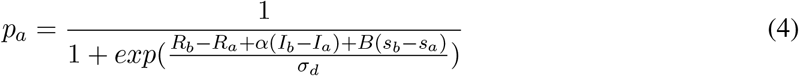

We simply define the information (I) in a way that when option b is more informative than a, *I*_*b*_−*I*_*a*_ = +1 and if option a is more informative than b, *I*_*b*_−*I*_*a*_ = −1 and if they are equal in information *I*_*b*_−*I*_*a*_ = 0. Similarly, the location variable is defined in a way that *s*_*a*_−*s*_*b*_ = +1 if option b is on the right and *s*_*a*_−*s*_*b*_ = 1 if it is on the left.

By fitting this model into our data, we are able to obtain information bonus (*α*), spatial bias (B) and decision noise (*σ*_*d*_) for each participant. We expect that information bonus (*α*) would be highly correlated with our model-free parameter p(high info) and decision noise (*σ*_*d*_) with p(low mean).

### 2- Model-based Results

Figure S1 shows the scatter plots between corresponding model-free and model-based parameters. As it was expected, we see a high correlation between information bonus and p(high info) and between decision noise and p(low mean) in both horizon. Model-based directed exploration (as defined by: information bonus h6 - information bonus h1) and model-based random exploration (decision noise h6 - decision noise h1) were also highly correlated with model-free directed and random explorations, respectively. Figure S2 shows the scatter plots and correlations between temporal discounting and the model-based parameters. The relationships are almost the same as the relations between temporal discounting and model-free parameters. Given this high correspondence, we based our main analyses in the manuscript on the model-free parameters.

**Figure S1:**
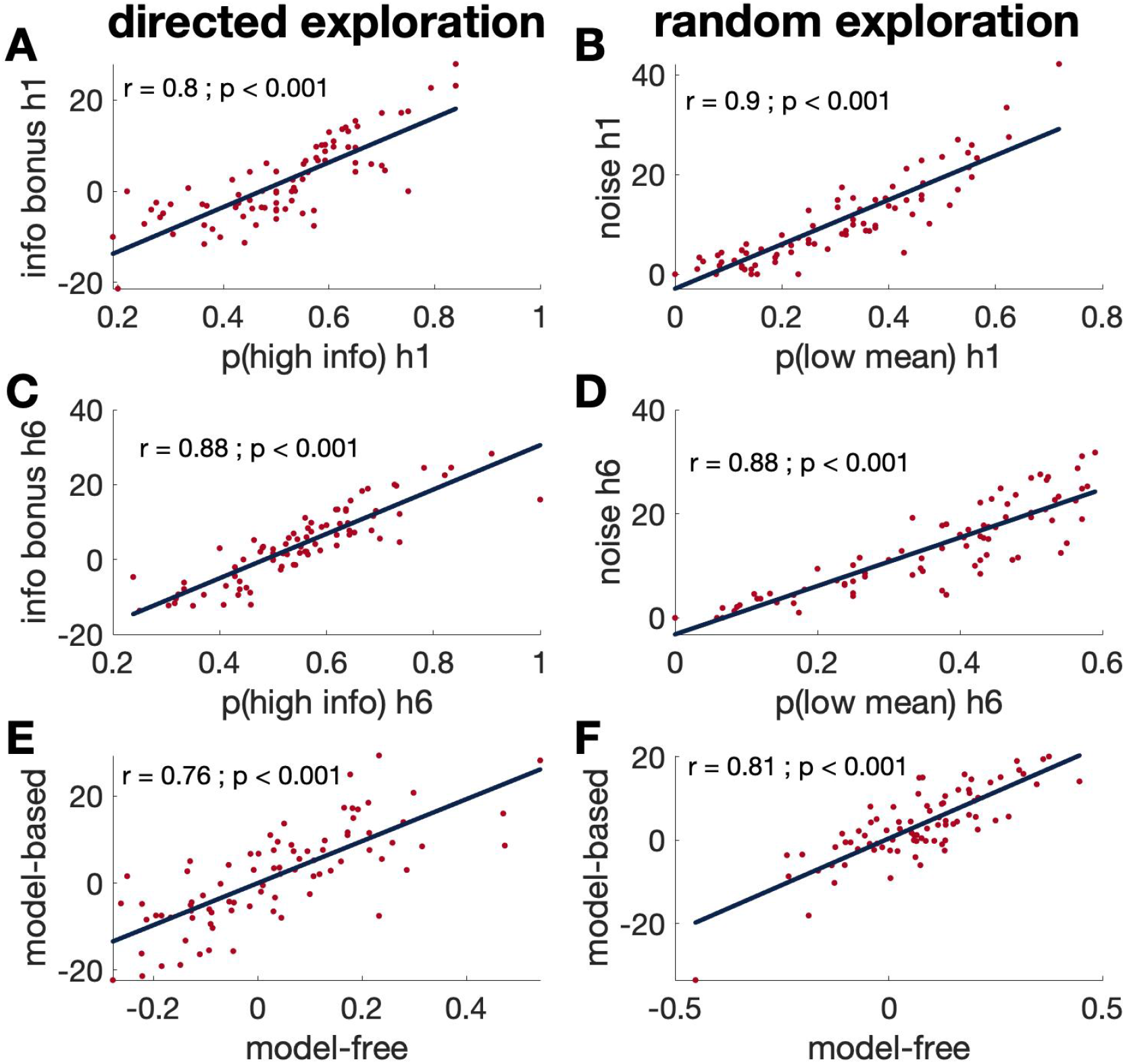
Scatterplots / Correlations between model-free and model-based parameters: (A) information bonus (from model) and p(high info) in horizon 1, (C) information bonus and p(high info) in horizon 6, (E) model-based directed exploration = information bonus h6 - information bonus h1, model-free directed exploration = p(high info) h6 - p(high info) h1, (B) decision noise (from model) and p(low mean) in horizon 1, (D) decision noise and p(low mean) in horizon 6, (F) model-based random exploration = decision noise h6 - decision noise h1, model-free random exploration = p(low mean) h6 - p(low mean) h1

**Figure S2:**
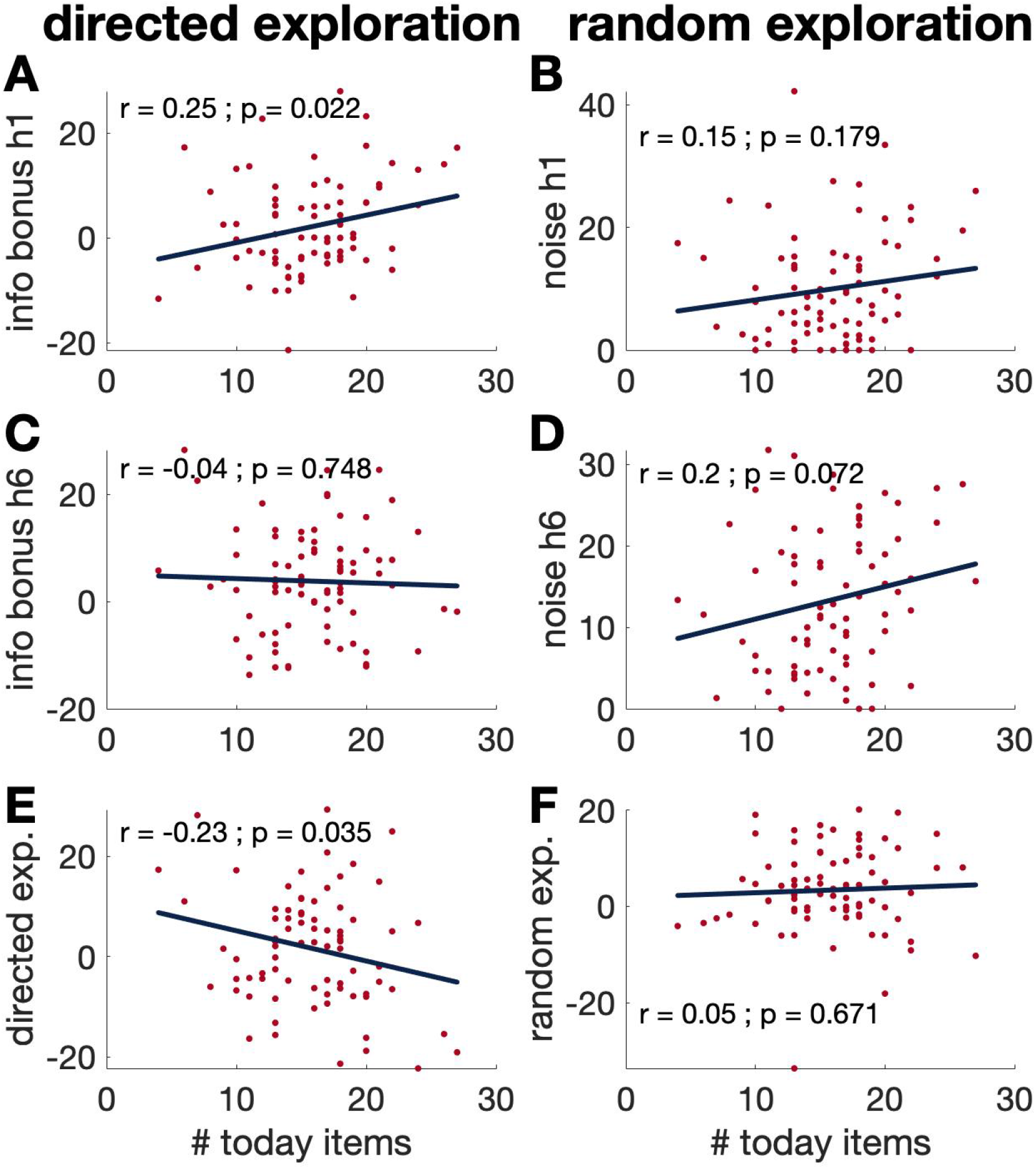
Scatter plots / correlations for model-based parameters over a temporal discounting measure (# today items). (A) information bonus at horizon 1, (B) decision noise at horizon 1, (C) information bonus at horizon 1, (D) information bonus at horizon 6, (E) model-based directed exploration (= info bonus h6 - info bonus h1, and (F) model-based random exploration (= noise h6 - noise h1. It yields the exact same conclusion we reached by a model-free analysis

### 3- Correlations between temporal discounting measures

Figure S3 shows that there is high correlations between different measures of temporal discounting in our study. More specifically, there is high correlations between a simple measure of temporal discounting, i.e. # today items: the total number of today (smaller immediate items) chosen by subject and more sophisticated measures (different k’s) that utilize a model fitting (Pearson’s correlation coefficients between .89 - 1). So we selected the simplest one (# today items) in our main analysis.

**Figure S3:**
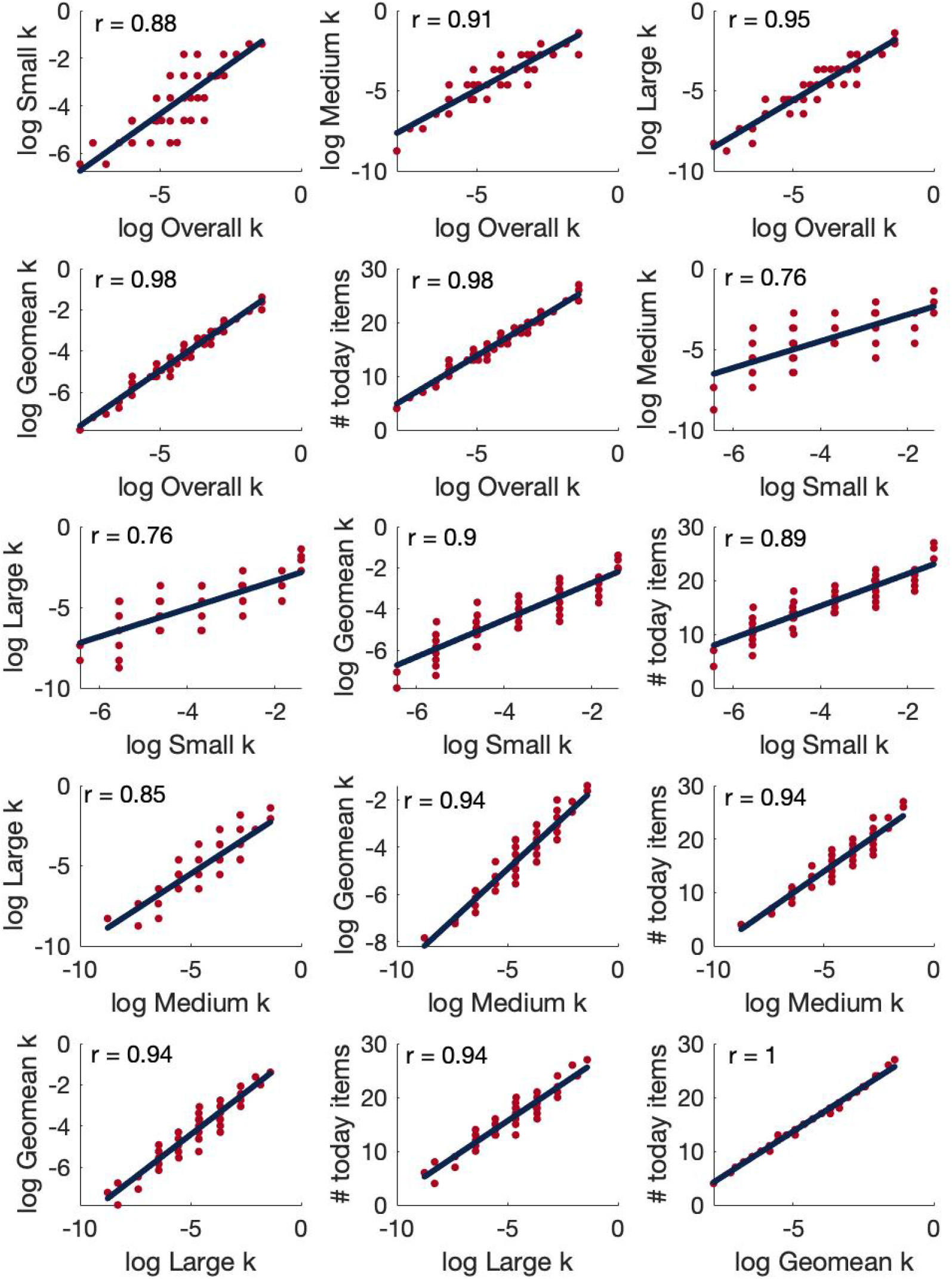
Scatterplots / Correlations between different measures of temporal discounting: log Overall k, log Small k, log Medium k, log Large k, log Geomean k, and # today items

### 4- Instructions for Horizon Task

Page 1: ‘Welcome! Thank you for volunteering for this experiment.’

Page 2: ‘In this experiment you will do four things.

1. Baseline eye measurement. This will take about 5 minutes.
2. Play a gambling task in which you will make choices between two options. This will take about 20 minutes.
3. Play another gambling task which also takes about 20 minutes.
4. When you”re done with the tasks, there will be a short post-experiment survey.’

Page 3: ‘Now we are going to get a baseline measurement of your eyes using the eye tracker.’

Page 4: ‘To do this we need you to stare at the screen for 5 minutes. Feel free to relax and daydream, but please stay in the chin rest. Press space to continue’

Page 5: ‘Press space to start the eye-measurement.’

After 5 minutes baseline eye tracker:

Page 1: ‘Welcome! Thank you for volunteering for this experiment.’

Page 2: ‘In this experiment - the gambling task - we would like you to choose between two one-armed bandits of the sort you might find in a casino.’

Page 3: ‘The one-armed bandits will be represented like this’

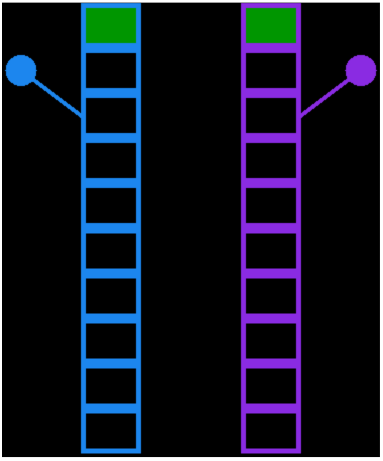

Page 4: ‘Every time you choose to play a particular bandit, the lever will be pulled like this …’

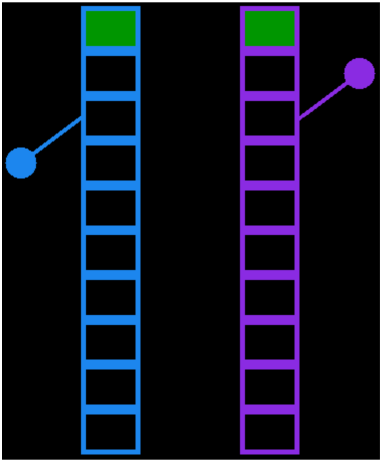

Page 5: ‘… and the payoff will be shown like this. For example, in this case, the left bandit has been played and is paying out 77 points.’

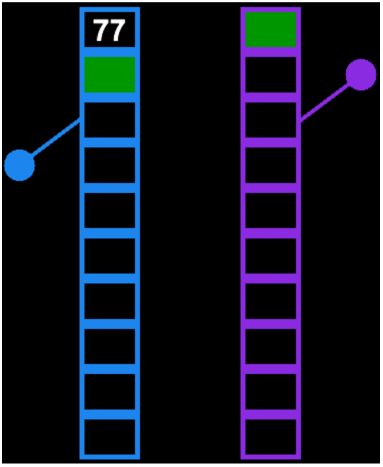

Page 6: ‘During one game, each bandit tends to pay out about the same amount of reward on average, but there is variability in the reward on any given play.’

Page 7: ‘For example, the average reward for the bandit on the right might be 50 points, but on the first play we might see a reward of 52 points because of the variability …’

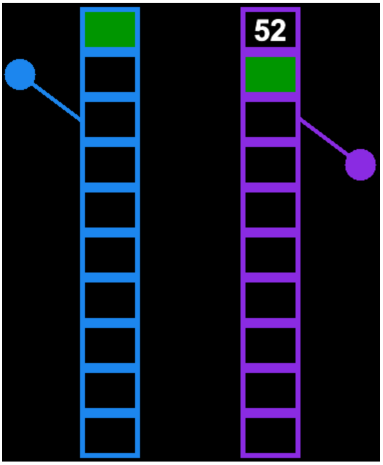

Page 8: ‘… on the second play we might get 56 points … ’

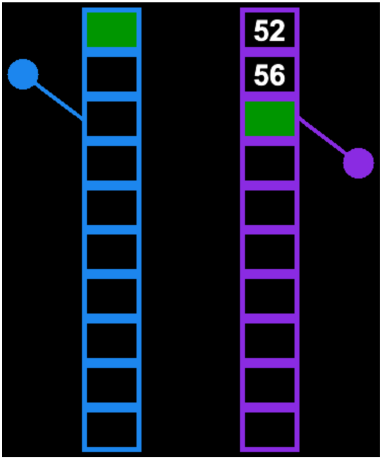

Page 9: ‘… if we open a third box on the right we might get 45 points this time … ’

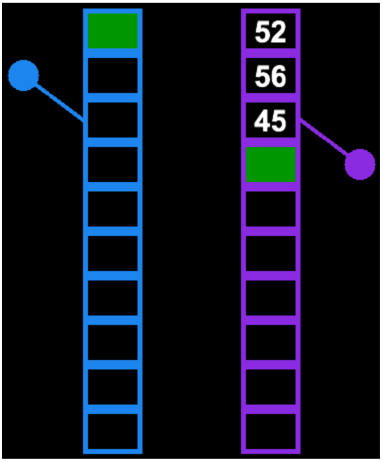

Page 10: ‘… and so on, such that if we were to play the right bandit 10 times in a row we might see these rewards …’

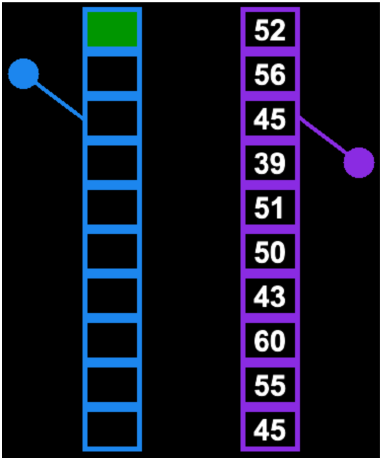

Page 11: ‘Both bandits will have the same kind of variability and this variability will stay constant throughout the experiment.’

Page 12: ‘During one game, one of the bandits will always have a higher average reward and hence is the better option to choose on average.’

Page 13: ‘To make your choice: Press <- to play the left bandit Press -> to play the right bandit’

Page 14: ‘On any trial you can only play one bandit and the number of trials in each game is determined by the height of the bandits. For example, when the bandits are 10 boxes high, there are 10 trials in each game … ’

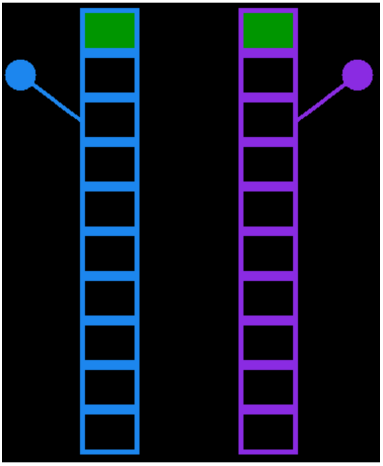

Page 15: ‘… when the stacks are 5 boxes high there are only 5 trials in the game.’

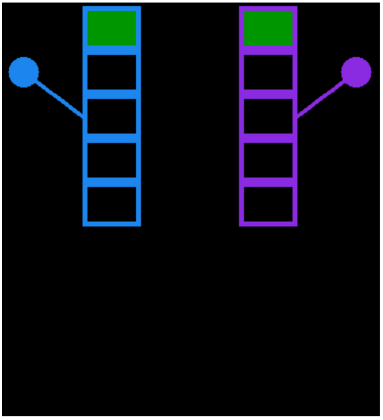

Page 16: ‘In addition, the first 4 choices in each game are instructed trials where we will choose an option for you. This will give you some experience with each option before you make your first choice.’

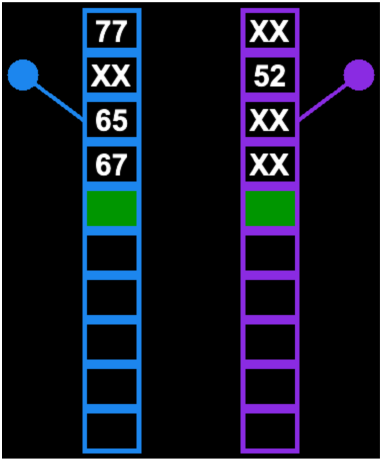

Page 17: ‘These instructed trials will be indicated by a green square inside the box we want you to open and you must press the button to choose this option in order to move on to see the reward and move on the next trial. For example, if you are instructed to choose the left box on the first trial, you will see this:’

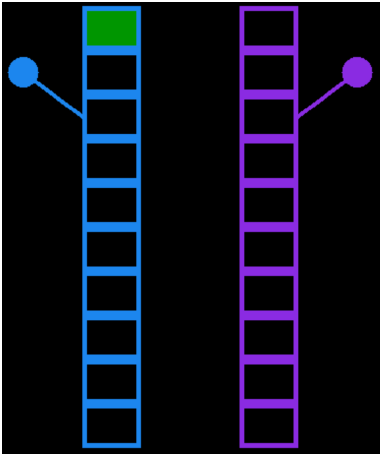

Page 18: ‘If you are instructed to choose the right box on the second trial, you will see this:’

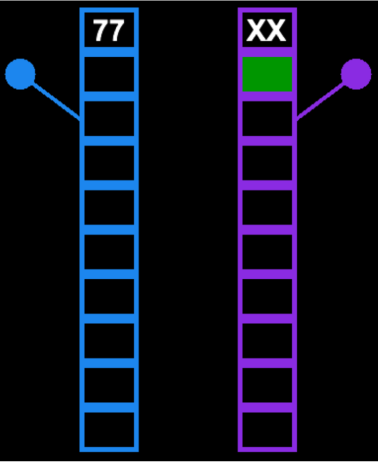

Page 19: ‘Once these instructed trials are complete there will be a go-cue and a beep, and then you will have a free choice between the two stacks that is indicated by two green squares inside the two boxes you are choosing between.’

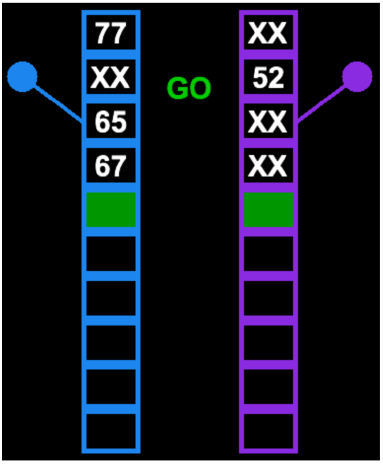

Page 20: ‘Throughout the task we will be tracking your eyes. To help us better track your eyes, the timing of the task is quite slow. Each game begins with the presentation of a fixation cross like this … please try to stare at this cross while it is displayed.’

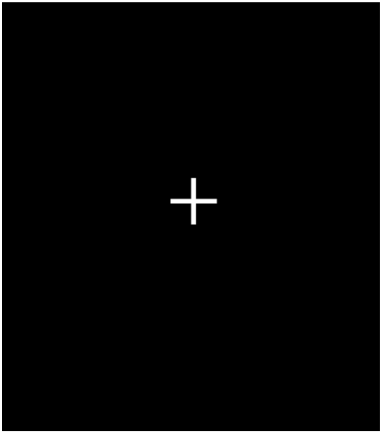

Page 21: ‘Then, the fixation cross will disappear and the bandits will appear - like this … during this period feel free to look where you want to.’

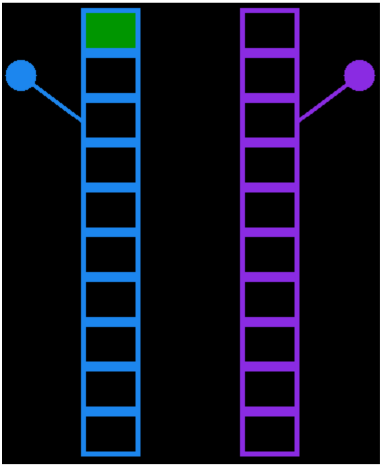

Page 22: ‘Then, a GO cue will appear at which point you need to choose between the two options’

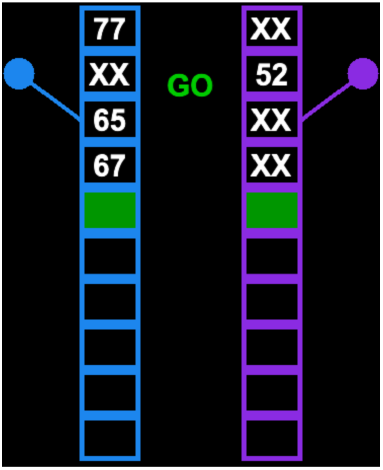

Page 23: ‘So … to be sure that everything makes sense let’s work through an example game … Press <- to play the left bandit Press -> to play the right bandit’

Page 25: ‘Good job! Now you know how to play this game.’

Page 26: ‘Press space when you are ready to begin. Earn as many points as you can! Good luck! Remember to stay in the chin rest!’

